# The prenatal exposome and genome in predictive modelling of DNA methylation

**DOI:** 10.64898/2026.08.12.742972

**Authors:** Rosa H. Mulder, Elena Isaevska, Claudio Cappadona, Serena Defina, Alexander Neumann, Janine F. Felix, Esther Walton, Matthew Suderman, Charlotte A. M. Cecil

## Abstract

**Introduction:** Fetal development represents a critical window during which genetic and environmental influences shape lifelong health. DNA methylation (DNAm) is a candidate underlying mechanism. While individual prenatal exposures have been related to DNAm, no studies have investigated the broader prenatal exposome, nor incorporated genetics with the exposome. Here, we integrated the prenatal exposome and genetics as predictors of DNAm at birth.

**Methods:** We used data from the Dutch Generation R (*n*=2282) and English Avon Longitudinal Study of Parents and Children (ALSPAC; *n*=809) cohorts. We performed epigenome-wide elastic net regression, using Generation R for model development/internal validation and ALSPAC for external validation, to predict DNAm at each CpG site. We used three models: Model 1 included 42 prenatal exposures, Model 2 additionally included child sex, gestational age and birth weight, and Model 3 further included meQTLs.

**Results:** In Model 1, the prenatal exposome explained on average 0.7% of DNAm variation across 347 validated CpGs (0.1% of tested CpGs). This increased to 40,044 CpGs (10.2%) with 1.3% of variation explained in Model 2, and 91,305 CpGs (23.2%) with 3.0% of variation explained in Model 3. In Model 1, prenatal smoking was the largest predictor, followed by delivery characteristics, among which meconium-stained amniotic fluid was a novel finding. In Model 3, typically both SNPs and multiple prenatal exposures were selected.

**Discussion:** We find that genomic associations with cord blood DNAm are stronger and more widespread than prenatal exposures, although typically, the prenatal exposome explains additional variation in DNAm beyond genetic influences.

## Introduction

Fetal development represents a critical window during which environmental influences can have lasting effects on health across the life course^1–3^. A wide range of prenatal exposures – such as suboptimal maternal nutrition or smoking during pregnancy – have been associated with adverse outcomes in childhood and adulthood^4,5^. Epigenetics is often considered a potential mechanism through which such prenatal factors might influence fetal growth and development^1,6^. In particular, DNA methylation (DNAm) – the addition of methyl groups to the DNA helix, generally at a cytosine-guanine (CpG) site – has been extensively studied in this context. Indeed, many epigenome-wide association studies have linked prenatal exposures to cord blood DNAm, with maternal smoking emerging as one of the most commonly investigated and robustly associated exposure^7^. Weaker associations with DNAm have also been reported for a range of other factors, such as maternal exposure to air pollution^8,9^, maternal BMI^10^, and maternal anxiety during pregnancy^11^.

Importantly these existing studies have focused on individual prenatal exposures in isolation. This is only partly informative, considering that different environmental factors are often intercorrelated^12^. For example, lower educational achievement is related to a range of risk factors such as poor nutrition, increased stress, and unhealthy living environment^13^. Lack of estimates taking this interdependence into account make it difficult to infer how much the environment contributes to wDNAm as whole based on these studies, thus studying the broader prenatal exposome rather than separate univariate exposures may give broader insights into how relevant the prenatal environment is to DNAm at birth.

To the best of our knowledge, studies on the combined impact of prenatal exposures on DNAm at birth have not yet been performed. Maitre et al.^14^ studied individual associations between over 100 prenatal exposures and DNAm in childhood in 1301 mother-child pairs and found that maternal smoking during pregnancy was a main predictor of DNAm. However, (i) exposures were studied successively and independently, thereby not taking into account multivariate contributions across the exposome. Moreover, (ii) childhood DNAm is influenced by both prenatal and postnatal exposures whereas cord blood DNAm at birth, which is investigated here, precedes postnatal exposures thereby enabling to better isolate the role of prenatal exposures. Lastly, (iii) it is known that there are widespread, strong influences of genetics on DNAm^15^, but how these compare to prenatal environmental exposures is not yet clear. Previously, we showed that genetic influences on DNAm at birth were much more frequently detected (by a 100,000 fold) than that of maternal prenatal stress^16^. Another study similarly found that genetic influences more often explained variation in DNAm at birth than prenatal exposures, but also that prenatal exposures often added explained variance on top of genetics^17^. However, studies that directly compare effects sizes of genetics with those of other prenatal factors are scarce and do not take the prenatal exposome into account as a whole, nor the combination of the exposome and the genome. Understanding the associations of the environment and the genome with DNAm can help to disentangle the extent to which epigenetic variation is explained by inherited versus potentially modifiable factors.

Here we used data from two large birth cohorts to address these gaps. The objectives of this study were to estimate the association between the prenatal exposome and child DNAm at birth, and to understand how these associations compare to those with the child’s genome. We did this by combining the methods of classical epigenome-wide analyses (EWAS) with elastic net methods to select relevant exposome-related features for each CpG – which were tested in one cohort and then validated in an independent one. We also ran elastic net models including SNPs on top of exposome-related features. We hypothesized that among all prenatal exposures, maternal prenatal smoking would be the largest contributor and that the genome would explain more variance in DNAm than the prenatal exposures.

## Methods

### Setting

In this study, data from the Generation R Study were used for development and internal validation of the models and data from the Avon Longitudinal Study of Children and Parents (ALSPAC) were used as an external validation sample. In the Generation R Study, pregnant women residing in the study area of Rotterdam in the Netherlands with an expected delivery date between April 2002 and January 2006 were invited to participate in the study^18^.

In ALSPAC, pregnant women resident in Avon, UK with expected dates of delivery between 1st April 1991 and 31st December 1992 were invited to take part in the study^19,20^. The ALSPAC website contains details of all the data that are available through a fully searchable data dictionary and variable search tool (http://www.bristol.ac.uk/alspac/researchers/our-data/).

### Ethics

The Generation R Study is conducted in accordance with the World Medical Association Declaration of Helsinki and has been approved by the Medical Ethics Committee of Erasmus MC, University Medical Center Rotterdam. Informed consent was obtained for all participants. Ethical approval for ALSPAC was obtained from the ALSPAC Ethics and Law Committee and the Local Research Ethics Committees. Consent for biological samples has been collected in accordance with the Human Tissue Act (2004). Informed consent for the use of data collected via questionnaires and clinics was obtained from participants following the recommendations of the ALSPAC Ethics and Law Committee at the time.

### Sample selection

In Generation R, the included pregnant mothers had 9,749 live-born children. DNA methylation was measured in cord blood of 1,393 children with the Illumina Infinium HumanMethylation450 BeadChip (Illumina Inc., San Diego, CA) (*GENR 450K*), and in cord blood of a second set of 1,115 children with the Infinium MethylationEPIC v1.0 Beadchip (*GENR EPIC*), resulting in a combined epigenetic dataset of 2,508 children. Genetic data were available for 2,444 of these children. Of these, 2,428 children had data on at least 50% of prenatal exposure variables as well (mean [SD] percentage of variables present 91% [11%]). Finally, participant pairs with up to second-degree relatedness were identified (IBD≥0.125) among the combined *GENR 450K* and *GENR EPIC* set and from each pair, one participant with the lowest number of prenatal exposure observations, or otherwise randomly, was excluded, resulting in a final sample of 2,282 (1,254 for *GENR 450K* and 1,028 for *GENR EPIC*) children, all of European ancestry^21^.

For a full overview of ALSPAC phases of enrollment, we refer to **Supplementary Information 1** and other publications^20,22^. In brief, the initial number of pregnancies enrolled was 14,541. Of these, 13,988 children were alive at 1 year of age. When the oldest children were approximately 7 years of age, an attempt was made to bolster the initial sample with eligible cases who had failed to join the study originally. The phases of enrolment are described in more detail elsewhere^20,22^. The total sample size for analyses using any data collected after the age of seven is therefore 15,447 pregnancies, resulting in 15,658 foetuses. Cord blood epigenetic data was available for a subsample of 912 of these children as part of the Accessible Resource for Integrated Epigenomic Studies (ARIES) study^23^. Of these, 842 children had genetic data available and of these, 817 had data on at least 50% of prenatal exposure variables (mean [SD] percentage of variables present 91% [7%]). Last, from each pair with up to second-degree relatedness (IBD≥0.125), one participant with the lowest number of prenatal exposure observations, or otherwise randomly, was excluded, resulting in a final sample with 809 children, all of European ancestry.

### Genetics

Overlapping SNPs between Generation R and ALSPAC were selected after cohort-specific quality control. In Generation R, children were genotyped with the Illumina HumanHap 610 or 660 quad chips. A full description has been published previously^21^. PLINK 1.90 was used for quality control^24^. Samples were excluded in the case of sex mismatches, minimal or excessive heterozygosity, or a sample call rate of <97.5%. Data were imputed to the 1000 genomes reference panel (Phase 1 version 3). Phasing was done using MACH software, and imputation using Minimac software. The ALSPAC children have been genotyped with the Illumina HumanHap 550 quad chip. The data were imputed to a phased version of the 1000 genomes references panel (Phase 1 version 3) from the Impute2 reference data repository. Best-guess genotypes were used in both cohorts. Autosomal variants were selected and variants with SNP call rates of <95%, with evidence for violation of Hardy-Weinberg equilibrium (p<1×10^-07^), with a minor allele frequency <5%, or with low imputation quality (Rsq<0.3 in Generation R and info scores <0.8 in ALSPAC, according to local practices^21,25^) were removed. Insertions, deletions, and multi-allelic positions were also removed. This resulted in 5,568,766 SNPs in Generation R and 5,886,328 SNPs in ALSPAC, of which 5,382,602 overlapped. To ensure consistent allele coding across cohorts, the effect allele was set to the minor allele based on the minor allele frequency in Generation R.

Second, SNPs identified as methylation quantitative trait loci (meQTLs), associated with DNAm were selected. MeQTL associations were identified using the largest available resources not including Generation R and ALSPAC, based on 171 cord blood samples (all *cis*-meQTLs)^26^ and on 4,170 peripheral blood samples of adults (*cis-* and *trans-* meQTLs)^27^. A total of 2,358,127 meQTLs were present in both the Generation R and ALSPAC genetic datasets (26% of CpGs had at least one meQTL [median = 56; range =1-4986]).

### Prenatal environmental exposures

All relevant data were obtained during pregnancy via questionnaire, at the research center or via medical or obstetric records. Data of Generation R and ALSPAC were compared and prenatal environmental exposures were selected based on availability in both cohorts and previously shown relevance to health and/or DNAm (**Supplementary Table 1**). Furthermore, variables with data availability in <50% of the cohort were excluded (no variables excluded) and categorical variables were only included if at least 30 participants in the selected samples were exposed (gestational diabetes and preeclampsia excluded). A list of all selected exposures is depicted in **Supplementary Table 1**, together with motivation for inclusion based on previous literature and a description of harmonization steps taken. This selection procedure resulted in a set of 42 prenatal exposure variables, spanning maternal characteristics (e.g. maternal age); maternal physical health (e.g. gestational hypertension and influenza infection); delivery characteristics (e.g. mode of delivery); maternal mental health (e.g. depressive symptoms during pregnancy); maternal environment (e.g. air pollution and distance to green spaces during pregnancy); and maternal lifestyle (e.g. smoking during pregnancy and nutritional factors). Missing datapoints (average (SD) missingness in final samples was 8% (5%) for Generation R and 6% (10%) for ALSPAC; ranges 0-21% and 0-37%, respectively **Supplementary Table 2**) were imputed in each cohort using *imputeFAMD* of the *missMDA* R package, employing a factorial analysis with a regularized iterative algorithm^28^.

### DNA methylation

For both cohorts, DNA extracted from cord blood was bisulfite converted. The EZ-96 DNAm kit (shallow) (Zymo Research Corporation, Irvine, CA) was used for bisulfite conversion. Samples were processed with the Illumina Infinium HumanMethylation450 BeadChip (Illumina Inc., San Diego, CA) in *GENR 450K* and *ALSPAC 450K* and with the Infinium MethylationEPIC v1.0 Beadchip in *GENR EPIC*.

In *GENR 450K* and *GENR EPIC*, the CPACOR workflow^29^ was applied for quality control. Arrays with observed technical problems such as failed bisulfite conversion, hybridization or extension as well as arrays with a sex mismatch were removed. Arrays with a call rate >95% in *GENR 450K* and >96% in *GENR EPIC* per sample were carried forward into normalization. In *ALSPAC 450K*, the meffil package in R version 3.4.3 was used for quality control^30,31^. Samples with mismatched genotypes, mismatched sex, incorrect relatedness, low concordance with samples collected at other time points, extreme dye bias and poor probe detection were removed and carried before normalization.

In the *GENR 450K and GENR EPIC*, data were quantile normalized in R^30^. In ALSPAC, functional normalization was performed (using 10 control probe principle components with slide included as a random effect) with the meffil package in R^31^. Probes were excluded if they had a detection *p*>0.01 or low bead count (<3) in >10% of the samples. DNAm levels were represented as beta values, indicating the ratio of methylated signal relative to the sum of methylated and unmethylated signal per CpG.

For the three (sub-)cohorts, 393,298 overlapping autosomal CpGs were present. To enhance precision of the analyses, the epigenetic sets of each cohort were residualized for batch (using a random intercept for 37 sample plates and for array [450K/EPIC] in Generation R and for 20 surrogate variables in ALSPAC), estimated white blood cell proportions, and 5 genetic PCs. The white blood cell proportions (CD4+ T-lymphocytes, CD8+ T-lymphocytes, natural killer cells, B-lymphocytes, monocytes, granulocytes, and nucleated red blood cells) of the cord blood samples was estimated using a cord blood reference panel^32^. After residualizing, beta values of each CpG outside the range of (25th percentile - 3*interquartile range (IQR)) to (75th percentile + 3*IQR) were winsorized to reduce the influence of extreme outlying values.

### Statistics

#### Epigenome-wide elastic net modelling

Associations between the prenatal exposome and DNAm were tested using elastic net regression with prenatal environmental exposures as features and each CpG as an iterative outcome. We used a two-stage validation design with ElasticNetCV from the scikit learn package in Python version 3^33^ (**Figure 1**). Eighty percent of the Generation R dataset (including both *GENR 450K* and *GENR EPIC* samples) was selected randomly as development sample using an internal 10-fold cross-validation (3 repeats) and hyperparameter tuning (net regularization strength [scikit-learn alpha, corresponding to in the glmnet notation]=[0.00001, 0.0001, 0.001, 0.01, 0.1, 1.0, 10, 100], L1/L2 mixing parameter [scikit-learn l1_ratio, corresponding to in the glmnet notation]=[0.1, 0.2, 0.4, 0.6, 0.8, 1.0]), the other 20% (including both *GENR 450K* and *GENR EPIC* samples) was used as an internal validation sample, and the ALSPAC dataset was then used as an independent external validation sample.

**Figure 1.**
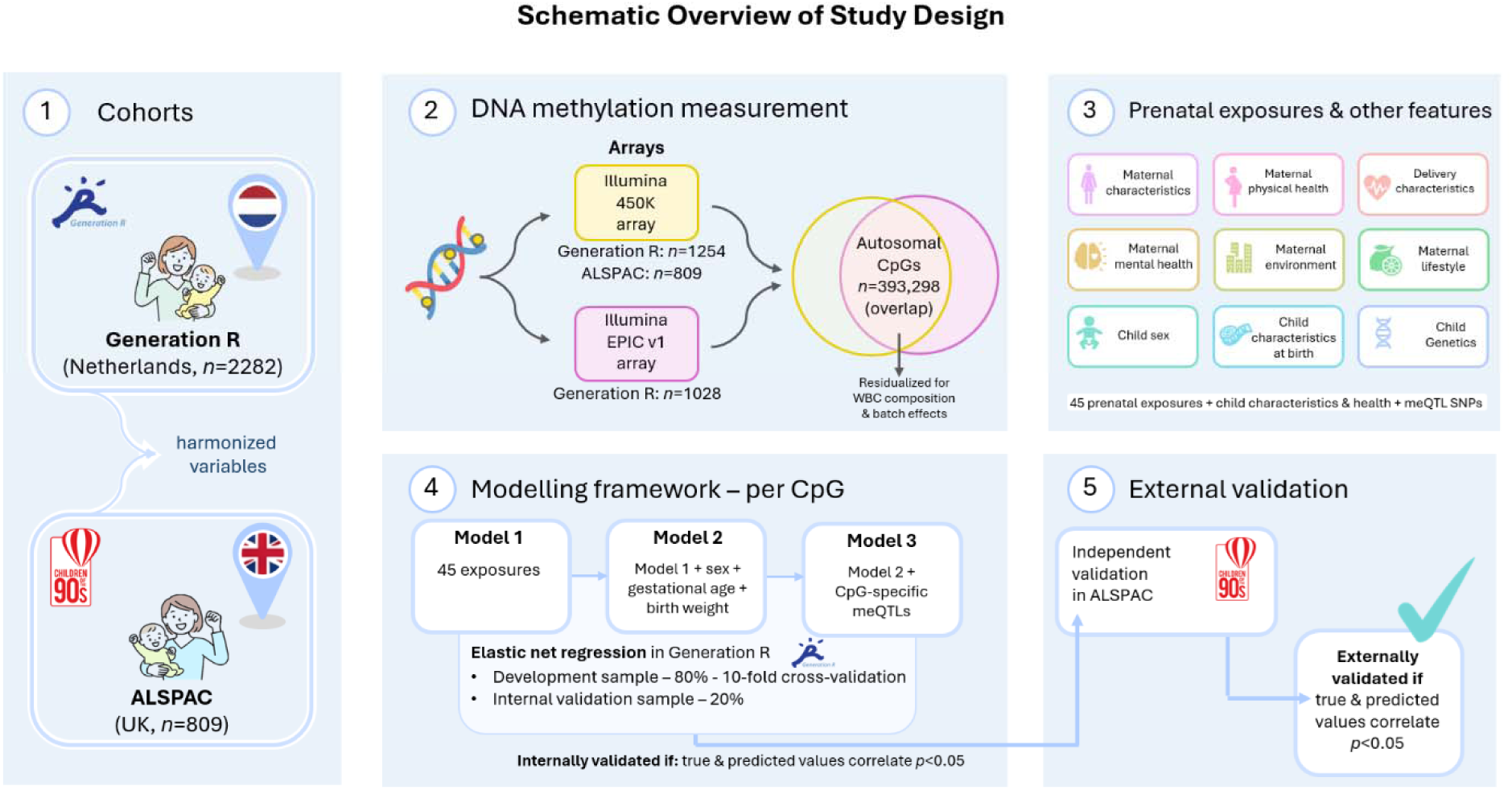
Study design

For each CpG, we performed three analyses to predict DNAm values residualized for technical covariates. In *Model 1*, only prenatal environmental exposures were included. In *Model 2*, we added child sex, gestational age and birth weight, i.e. variables that are often included as covariates in EWASs. Last, in *Model 3*, we additionally included known meQTLs (specific to each CpG) for the CpGs that had at least one meQTL. We set the maximum number of included meQTLs to 50, to have a comparable number of prenatal exposures and meQTLs. If a CpG had more than 50 meQTLs, feature selection using univariate linear regression (*f_regression*) was performed to select the 50 most strongly associated meQTLs. Categorical variables that had more than two levels were divided into dummy variables, hence elastic net regression was performed on a total of 45 features in Model 1, 48 features in Model 2, and a maximum of 98 features in Model 3. Values for the features and CpGs in the three samples (development-, internal validation-, external validation sample) were scaled and set to a value between 0 and 1. As a performance criterion, we used *p*-values, which is common in epigenome-wide analyses, over R^2^, which is more common in machine learning paradigms – since this is the first study of its kind, we were not able to form expectations of how much explained variance would be meaningful. CpGs were considered internally validated if the predicted and true values in the internal validation sample had a positive Pearson correlation with *p*<0.05. CpGs were considered to be externally validated if Pearson correlation between predicted and true values in both the internal validation and external validation sample had a positive Pearson correlation with *p*<0.05.

#### Results interpretation

We first evaluated predictive performance by reporting the number and percentage of CpGs for which prediction models were internally and externally validated. Analyses were restricted to externally validated CpGs. For these CpGs, we reported the median explained variance (R²) in the external validation cohort and the median number of predictors retained in the elastic net models. Differences between models in the proportion of externally validated CpGs were assessed using Fisher’s exact tests, whereas differences in explained variance among externally validated CpGs were assessed using unpaired T-tests (*p*<0.05). To contextualize our significance-based validation criterion, we additionally summarized the explained variance (R²) corresponding to validated models in both the internal and external validation samples and quantified the correlation of model performance (Pearson’s *r*) between cohorts across CpGs.

Second, we calculated the epigenome-wide cumulative coefficient of each predictor within a model. Because all predictors were standardized before model fitting, the absolute regression coefficients are directly comparable within each model. For each predictor, we therefore calculated its cumulative standardized coefficient by summing the absolute regression coefficients across all externally validated CpGs for which the predictor was retained. This epigenome-wide summary metric reflects both how frequently a predictor was selected across CpGs and the magnitude of its contribution when selected. We visualized these cumulative absolute coefficients in bar plots and used them to compare the overall contribution of predictors within and between models. To further disentangle differences between cumulative coefficients, we separately evaluated (i) differences in predictor selection frequency using Fisher’s exact tests and (ii) differences in coefficient magnitude among selected predictors using unpaired T-tests (*p*<0.05).

Finally, to evaluate the effect of adding covariates and genetic information to successive models, we compared the standardized regression coefficients of prenatal exposures between consecutive models. Analyses were restricted to exposure–CpG pairs for which the exposure was retained in both models, and differences in coefficient magnitude were assessed using paired T-tests (*p*<0.05).

### Sensitivity analysis

To assess the robustness of model performance to the random partitioning of the development and internal validation samples, we generated four random 80/20 splits of the Generation R cohort. Each CpG was randomly assigned to one of these splits, which was applied consistently across all three models. We then compared the proportion of externally validated CpGs across the four random splits using Chi-square tests (*p*<0.05).

### Follow-up analyses

Last, to understand if exposome-associated CpGs are more often related to later health and ageing, we performed two follow-up analyses on validated Model 1 CpGs. Firstly, we tested whether they show enrichment for CpGs suggestively related (*p*<1×10^-05^) to health outcomes in EWAS meta-analyses on cord blood DNAm. We used the results from five different Pregnancy and Childhood Epigenetics Consortium (PACE)^34^ EWASs on attention-deficit hyperactivity disorder symptoms (ADHD)^35^, general psychopathology^36^, asthma^37^, body mass index (BMI)^38^, and sleep duration^39^ - comprising a total of 162 suggestive CpGs – all harmonized in a larger meta-regression study^40^ (**Supplementary Table 3**). Fisher’s exact test was used to test if exposome-related CpGs were more often related to health outcomes than other CpGs were (*p*<0.05).

Secondly, we tested the enrichment of CpGs that have been selected into adult epigenetic clocks. We did this separately for first-generation clocks, which have been trained to estimate age, and for second- or third generation clocks, i.e. clocks trained to estimate mortality, time-to-death, or indicators of mortality. The inclusion and exclusion criteria for epigenetic clocks are described in **Supplementary Information 2**, which resulted in the selection of four first-generation clocks: Horvath’s clock,^41^ Hannum’s clock,^42^ Weidner’s (non-minimised) clock,^43^ Zhang’s Elastic Net clock^44^; nine second-generation clocks: PhenoAge,^45^ Huan’s ‘all-cause mortality and CVD death’ clock and ‘cancer death’ clock^46^, GrimAge^47^, Ying’s CausAge, DamAge, and AdaptAge^48^, DNAmTL^49^, and epiTOC2^50^; and a third-generation clock DunedinPACE,^51^ i.e. a clock trained on *change* estimates of age biomarkers to approximate pace of aging (**Supplementary Table 4**). Since second- and third-generation clocks are both trained on (indicators of) age-related diseases or mortality, instead of chronological age itself, these clocks were grouped together. Altogether, we included a total of 928 unique first-generation clock sites and 3974 unique second- and third-generation clock sites. Fisher’s exact test was used to test if exposome-related CpGs were more often selected into epigenetic clocks than other CpGs were (*p*<0.05).

## Results

Extensive descriptive statistics of the prenatal exposures, spanning domains of (i) maternal characteristics, (ii) maternal physical health, (ii) delivery characteristics, (iv) mental health, (v) environment, and (vi) lifestyle, as well as (vii) child sex and child (viii) characteristics at birth, are presented in **Supplementary Table 2** and histograms of the prenatal exposures are presented in **Supplementary Figure 1-2**. In brief, mothers were on average (SD) 32.0 (4.3) years of age when they gave birth in Generation R, and 29.7 (4.4) years in ALSPAC. The mothers were relatively healthy, with an average (SD) BMI of 23.1 (3.5) in Generation R, and 22.9 (3.7) in ALSPAC, and predominantly presented without anaemia during pregnancy (95% in Generation R and 92% in ALSPAC), and without gestational hypertension (95% in Generation R and 85% in ALSPAC). Most deliveries were spontaneous rather than assisted or via caesarian section (73% in Generation R and 74% in ALSPAC). Most mothers had completed a university degree in Generation R (62%) or university degree or A-levels in ALSPAC (49%) and most mothers did not smoke tobacco during pregnancy (77% in Generation R and 86% in ALSPAC) and most mothers in ALSPAC (66%) and nearly half of the mothers in Generation R (46%) did not drink alcohol during pregnancy. Of the children, 50% of the Generation R sample was female, compared to 51% in ALSPAC. Gestational age was on average (SD) 40.1 (1.5) weeks in Generation R and 39.6 (1.5) weeks in ALSPAC. Correlations between the predictors were abundant, and most strongly and consistently present in both cohorts between maternal depression, anxiety and stress during pregnancy, between the two air pollution measurements, between the food intake variables, and between gestational age at birth and birth weight (**Supplementary Figure 3-4**).

### Model performance

**Table 1** depicts the number of CpGs for which a model could be internally and externally validated. For Model 1, models for 347 CpGs (0.09%) were externally validated. For Model 2, this increased to 40,044 CpGs (10.18%), and for Model 3, 91,305 CpGs (23.22%). Increases in model performance (i.e. validation) were strongly statistically significant (Model 1 vs Model 2: OR=128.4, 95% CI=115.7-143.0, *p*<9.88×10^-324^; Model 2 vs Model 3: OR=2.7, 95% CI=2.6-2.7, *p*<9.88×10^-324^).

**Table 1.** Elastic net results for the different models.

|  | Internally validated<br>CpGs (n, %) | Internally & externally<br>validated CpGs (n, %) | Replication rate<br>(%) |
| --- | --- | --- | --- |
| Model 1 | 4494 (1.14) | 347 (0.09) | 7.7 |
| Model 2 | 63711 (16.20) | 40044 (10.18) | 62.9 |
| Model 3 | 120100 (30.54) | 91305 (23.22) | 76.0 |

Percentages were calculated relative to the total number of CpGs analyzed ( $n = 393,298$ ), not conditional on significance in the test set. Significance: $p < 0.05$ .
Model 1: included 42 exposures (dichotomized to 45 features)
Model 2: Model 1 + child sex + gestational age at birth and birth weight
Model 3: Model 2 + CpG-specific meQTLs

Additionally, the average amount of variance per CpG that could be explained by the validated models increased with each incremental model, with on average 0.7% (SD=1.3) of variance explained in Model 1, 1.3% (SD=2.7) in Model 2, and 3.0% (SD=15.6) in Model 3 (Model 1 vs Model 2: *t*(364.88)=-19.67, *p*=3.04×10^-59^; Model 2 vs Model 3: *t*(125598)=-157.65, *p*<9.88×10^-324^) (**Table 2**).

**Table 2.** Percentage of variance explained in externally replicated CpGs.

|  | Median (SD) |
| --- | --- |
| Model 1 | 0.7 (1.3) |
| Model 2 | 1.3 (2.7) |
| Model 3 | 3.0 (15.6) |

Model 1: included 42 exposures (dichotomized to 45 features); externally replicated 320 CpGs
Model 2: Model 1 + child sex + gestational age at birth and birth weight; externally replicated 4613 CpGs
Model 3: Model 2 + CpG-specific meQTLs; externally replicated 66492 CpGs

We used a threshold of *p*<0.05 for internal and external validation as a model performance criterion and inspected variance explained in the internal and external validation samples by the models established in the development sample. For each of the models (**Supplementary Figure 5**), *p*<0.05 corresponded to a minimum explained variance of R^2^=0.008% (*r*=0.09) in the internal validation sample, and R^2^=0.005% (*r*=0.07) in the external validation sample. Second, the correlation of explained variance for the CpGs, between the internal and external validation sample increased with each incremental model, with *r*=0.06 (*p*=1.25×10^-99^) in Model 1, *r*=0.58 (*p*<9.88×10^-324^) in Model 2, and *r*=0.80 (*p*<9.88×10^-324^) in Model 3.

### Epigenome-wide cumulative coefficient of predictors across models

We studied the epigenome-wide cumulative coefficient of each exposure within each model, by summing standardized absolute coefficients of each exposure over all externally validated CpGs. As such, the epigenome-wide cumulative coefficient indicates the total contribution of each exposure to the methylome (**Figure 2**; **Supplementary Table 5**). Among validated CpGs in Model 1, a median of 5 (SD=5.6; range=1-44) out of a total of 45 features were selected in the models. Only the size of nearest green space area was not retained as a predictor in any of the validated models. Of all features, continued prenatal smoking during pregnancy was the strongest contributor in Model 1. Contributions of delivery characteristics were also relatively high, with elective caesarian delivery as second highest, followed closely by meconium-stained amniotic fluid and assisted delivery. More specifically, the epigenome-wide cumulative coefficient of prenatal smoking was 10.7 times higher than the average epigenome-wide cumulative coefficient of all other Model 1 exposures. This was because continued prenatal smoking was more often selected as a contributor in the model (for 68.3% of validated CpGs) than the other exposures on average were (13.4%; OR=14.0, 95% CI=11.0-17.8; *p*=1.07×10^-117^), and because when selected into the model, the absolute coefficient was on average 2.4 times larger for continued prenatal smoking (M=0.02; SD=0.03), than it was for the other exposures (M=0.01; SD=0.01; t(244.02)=6.5, *p*=3.52×10^-10^).

**Figure 2.**
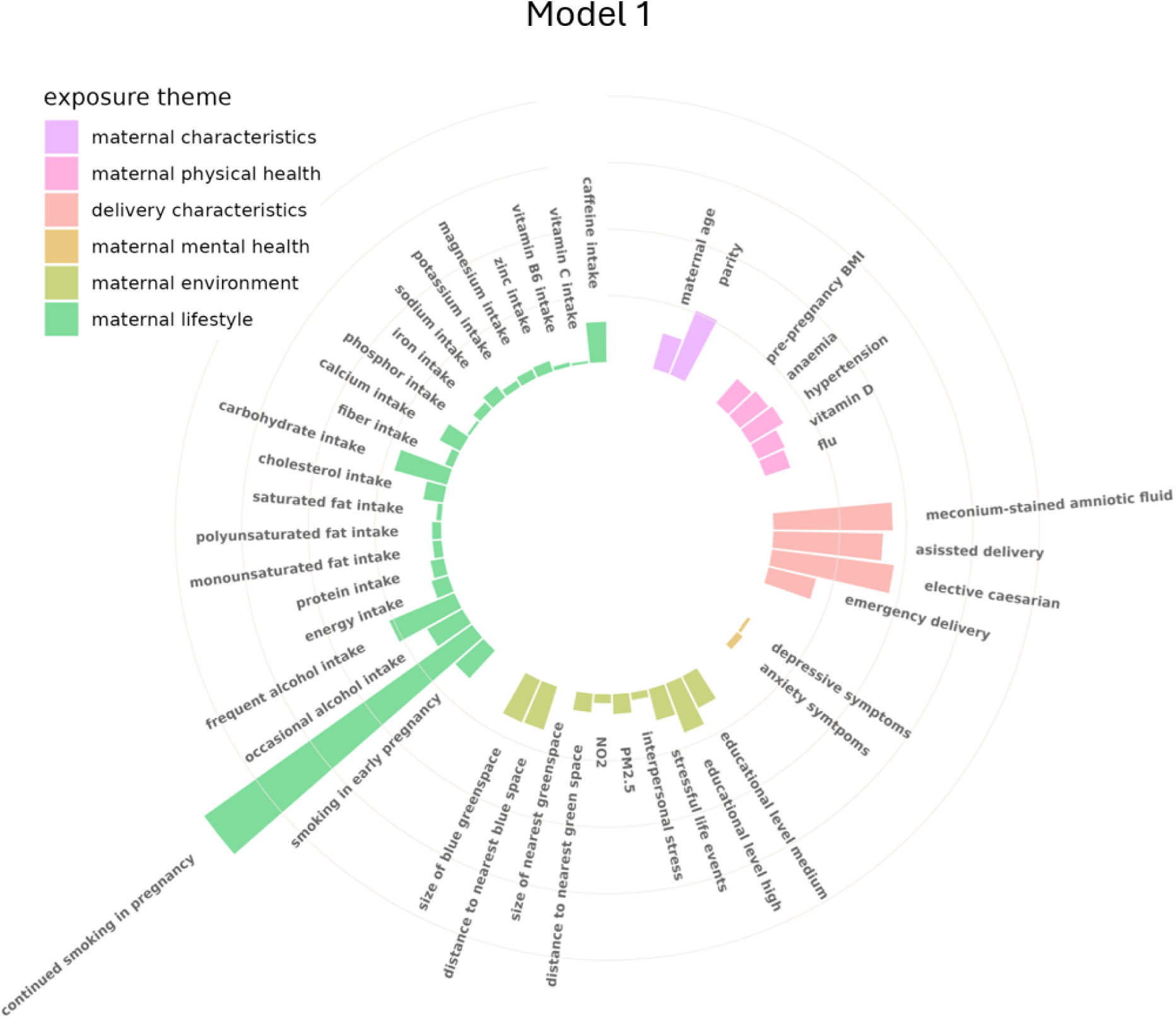
Epigenome-wide cumulative coefficient of exposures in Model 1. The length of the line represents the summed absolute regression coefficients across all externally validated CpGs. All features included in Model 1 are presented in the figure.

For Model 2, a median of 5 (SD=5.7; range=1-48) out of a total of 48 features were retained in the models. The epigenome-wide cumulative coefficient for the added predictors in Model 2, i.e. child sex, birth weight and gestational age, were respectively 11.7, 1.0, and 10.1 times higher than that of the largest contributor to Model 1, which was continued prenatal smoking (**Figure 3**; **Supplementary Table 6**). Hence child sex and gestational age added a relatively large proportion of explained variance in DNAm. Gestational age at birth was less often selected as an feature to Model 2 (31.9%) than prenatal smoking (38.7%; OR=0.7; 95% CI=0.7-0.8; *p*=1.19×10^-88^), but its average absolute coefficient (M=0.11; SD=0.08) was 14.2 times larger than that of prenatal smoking (M=0.01; SD=0.01; t(12980)= 154.7; *p*<9.88×10^-324^).

**Figure 3.**
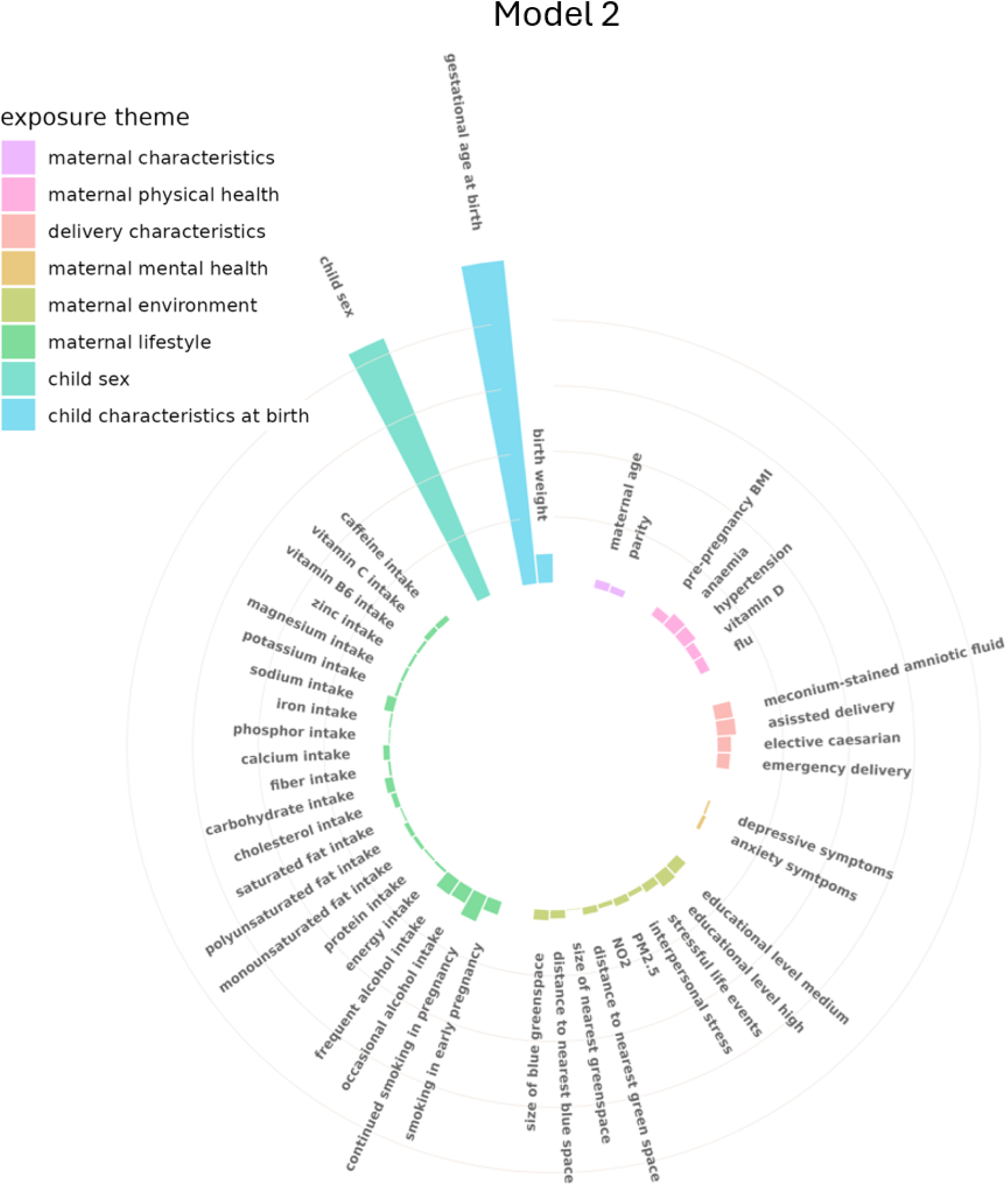
Epigenome-wide cumulative coefficient of predictors in Model 2. The length of the line represents the summed absolute regression coefficients across all externally validated CpGs. All features included in Model 2 are presented in the figure.

**Figure 4.**
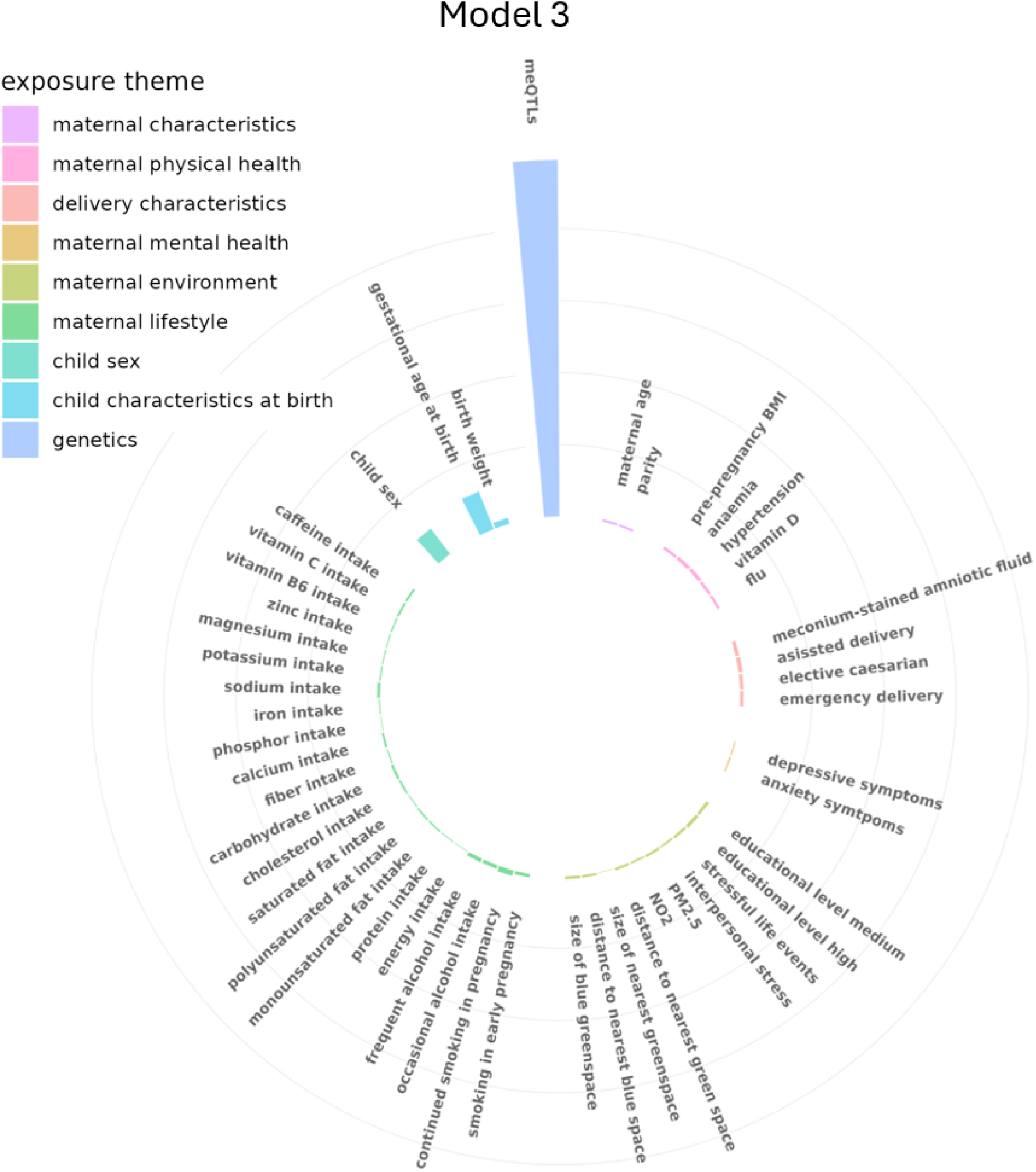
Epigenome-wide cumulative coefficient of predictors in Model 3. The length of the line represents the summed absolute regression coefficients across all externally validated CpGs. All features included in Model 3 are presented in the figure.

Last, in Model 3, a median of 13 (SD=13.6; range=1-98) out of 98 features were included in the validated models. A genetic factor was computed by adding the coefficients of all the different meQTLs into a single factor. Comparing the epigenome-wide cumulative coefficients among all predictors (**Figure 3**; **Supplementary Table 7-8**) showed this genetic factor has 8.8 times the weight of the next largest contributor, gestational age, and 64.8 times the weight of continued maternal smoking during pregnancy. When combining all features (exposures) from Model 1 together into one exposome factor, the genetic factor was 3.4 times larger. In most cases, the model included at least one meQTL (76.0%) and at least one exposure (i.e. features that were also included in Model 1; 89.7%) – yet the average number of meQTLs (M=8.5; SD=9.0; range 0-50 out of 50 features) was 1.2 times higher than the average number of exposures (M=6.9; SD=6.9; range 0-45 out of 45 features; t(91305)= -51.4; *p*<9.88×10^-324^). On top of that, the average absolute coefficient of the meQTLs selected into the models (M=4.3×10^-02^; SD=0.02) was 1.5 times higher than that of the selected exposures (M=2.8×10^-02^; SD=0.47; t(4127002)=6.5, *p*=8.01×10^-11^).

### Exposure level coefficients

To evaluate the effect of adding covariates and genetic information to successive models on predictor coefficients, we compared the coefficients of each exposure for the validated CpGs between the different models using paired T-tests. Comparing Model 1 to Model 2, we find indication that addition of child sex, gestational age and birth weight as predictors changed the size of the absolute coefficients for 11 of the 45 Model 1 features (*p*<0.05), with the average coefficient size decreasing for 5 features (meconium-stained fluid: -49%; assisted delivery: -18%; elective caesarian delivery: -29%; emergency caesarian delivery -18%; frequent alcohol intake: -22%) and increasing for 6 features (anaemia: 10%; hypertension: 14%; medium education level: 16%; smoking until pregnancy was known: 29%; continued smoking: 10%; occasional alcohol intake: 16%; full results are depicted in **Supplementary Table 9**). When comparing Model 2 to Model 3, we note that the size of the coefficient of 39 of the 48 features in Model 2 changed, with the average absolute estimate of 36 of the features decreasing after the addition of meQTLs to the model (mean [SD] change -7% [4%]), and increasing for 3 of the features (continued smoking: 2%; child sex: 2%; gestational age: 4%; full results are depicted in **Supplementary Table 10**), thereby showing that when taking genetics into account, the predictive value of prenatal exposures by and large seems to decrease.

### Sensitivity analysis

To assess the robustness of model performance to the random partitioning of the development and internal validation sample, we compared model performance between the different 80/20 splits that were performed. No difference could be detected in amount of validated models between the different splits, in any of the three models (all X^2^(393297)=262534, *p=*1.00).

### Follow-up analyses on health outcomes and ageing

To investigate potential implications of our findings for health outcomes and aging, we first tested if 347 validated Model 1 CpGs were enriched among suggestive findings (*p*<1×10^-05^) in cord blood EWASs of child health outcomes (*n*=159 CpGs). Among the validated Model 1 CpGs, no CpG was suggestively related to a child health outcome, hence there was no indication of enrichment. Second, we tested the enrichment of the appearance of validated Model 1 CpGs in first- or second- and third-generation clocks (*n*=928 CpGs and *n*=3974 CpGs, respectively). Among the validated Model 1 CpGs, 5 (1.4%) have been selected into first-generation clocks, which was relatively higher than the number of non-validated Model 1 CpGs (0.2%; OR=6.2; 95% CI=2.0-14.7; *p*=1.51×10^-03^). Furthermore, 11 validated Model 1 CpGs (3.2%) have been selected into second- or third generation clocks, which was relatively higher than that of non-validated Model 1 CpGs (1%; OR=3.2; 95% CI=1.6-5.8, *p*=9.53×10^-04^).

## Discussion

In this study, we investigated to what extent the prenatal exposome is related to child DNAm at birth using a hybrid epigenome-wide analysis and elastic net approach in two independent population-based cohorts. Leveraging harmonized data across 42 prenatal exposure variables (dichotomized into 45 features), we found that we could detect associations with exposures in DNAm at a minority of CpGs (n = 347; 0.1% of tested sites), and that the prenatal exposome on average explained a very small amount of variation per CpG (0.7%). When including child sex, gestational age at birth and birth weight to the model, this number increased to 40,044 CpGs (10.2%) with 1.3% of variation explained. When additionally taking into account child genetics by including known meQTLs, this number drastically increased to 91,305 CpGs (23.2%), with 3.0% of variation explained. Together, this indicates that genetics seem to repeatedly explain more variation in DNAm, with fewer and weaker associations between the prenatal exposome and DNAm. This finding is in line with our earlier study on genomic and prenatal stress associations with DNAm at birth^16^. However, we also found that most often a combination of both SNPs and prenatal exposures were selected as features in the model, indicating that, while the genome seems to have a stronger influence on DNAm, the prenatal exposome adds to this explained variance. Importantly, most of the validated models included multiple exposures, corroborating the idea that studying the prenatal exposome as a whole instead of univariate exposures is a better reflection of the real-life situation.

First, in line with our hypothesis and a previous large study on prenatal exposures and DNAm in childhood^14^, we found that among the prenatal exposures, maternal smoking during pregnancy was the largest contributor to DNAm variation at birth. Second, delivery characteristics seemed to have a relatively large contribution to the explained variance in DNAm at birth, with presence of meconium in the amniotic fluid as a notable variable that to the best of our knowledge has not yet been reported on. Meconium-stained amniotic fluid is a sign of fetal distress^52,53^. These associations might indicate a quick response of DNAm to fetal distress. On the other hand, we also found that elective caesarian delivery, which should not include fetal distress, was a relatively large contributor to DNAm, a finding that fits a recent report that caesarian delivery (elective and emergency caesarian delivery combined) is associated to DNAm at birth^54^. Since results showed that the inclusion of gestational age, birth weight and child sex tended to decrease the coefficients of the delivery characteristics, it might be that underlying conditions of the child or other pregnancy complications play a role in these associations. More research, potentially using mediation strategies or Mendelian randomization methods, could elucidate these pathways further.

In our follow-up analyses, we could not detect any enrichment for health outcome-related associations among exposome-related CpGs. However, this analysis might have been underpowered due to the low number of CpGs included in the enrichment analysis and focused on a limited set of childhood phenotypes for which multi-cohort EWAS statistics in cord blood are available. The follow-up analyses did show enrichment among exposome-related CpGs for both first- and second-/third-generation adult clock sites, indicating that CpGs that are related to the prenatal exposome, are also more often related to ageing or age-related health outcomes later in life. We previously showed that variation at birth in epigenetic clock sites is highly predictive of variation in early adulthood^55^, suggesting that related health associations later in life might already be determined early on.

These results should be interpreted in light of several considerations and limitations. First, the pre-selected meQTLs that were included to study the influence of genomics on DNAm, were based on resources that did not rely on ALSPAC and/or Generation R data^15,25^ – which meant that these resources were not the largest studies available. Yet, even with this limitation, genomics seemed to robustly explain a much larger part of DNAm variation than the prenatal exposome did. Second, we point out that the number of exposures we could test was much smaller than the number of meQTLs that have been identified, making it an unequal comparison between the two. With the exposome, we included factors that could be relevant and that we could harmonize between the two cohorts, but there are many factors that we were not able to include, such as other lifestyle exposures, chemical exposures, maternal metabolomics and proteomics^14^. However, we were able to span six different domains of exposures with variables that were present and harmonizable between two cohorts in different countries. Third, including gene-by-environmental interactions might have explained more variance in DNAm. However, in a previous study on prenatal stress, genetics and DNAm at birth, genetic main effects were much stronger and more prevalent than genetic-by-prenatal stress interaction effects^16^. While this may suggest a limited role of gene-by-environment interaction, testing more environmental features and the use of alternative methods like random forest models could prove to be more fruitful method to include interaction effects without dramatically increasing the number of features. However, the results would be less readily interpretable, as exposure coefficients are not outputted. Fourth, while we used two different populations in this study, both were of European ancestry. Research in a more diverse sample is necessary to understand to what extent our conclusions hold for other populations. Last, we would like to emphasize that prediction is not equal to causation. We therefore do not know if the prenatal exposures examined are causally related to DNAm, or how they are causally related to one another. Causal research, for example intervention studies for prenatal exposures or Mendelian Randomization studies could be applied to understand to what extent the prenatal exposome affects DNAm. Last, studies including long-term health outcomes are necessary to fully elucidate the role DNAm plays in relation to the prenatal environment and child health.

In conclusion, this is the first study to quantify the relative contribution of a range of prenatal exposures to DNAm patterns at birth. We find that prenatal smoking, as hypothesized, has the largest association with DNAm and also report for the first time a relatively large association of meconium-stained amniotic fluid. These associations are consistent across sets including different DNAm arrays, normalization methods and across two countries. Importantly, we see that the associations of prenatal exposures are dwarfed when taking into account the child genome, but that most often, the prenatal exposome and the genome both add to explaining variation in DNAm at birth.

## Contributors

RHM was involved in the conceptualization, methodology, formal analysis, visualization, writing the original draft and review and editing. EI was involved in reviewing and editing the manuscript. CC was involved in the formal analysis and reviewing and editing the manuscript. SD was involved in the methodology and reviewing and editing the manuscript. AN was involved in the methodology and reviewing and editing the manuscript. JFF was involved in data curation and reviewing and editing the manuscript. EW was involved in reviewing and editing the manuscript. MS was involved in reviewing and editing the manuscript. CAMC was involved in the conceptualization, methodology, reviewing and editing the manuscript and in supervision.

## Acknowledgements

The Generation R Study is conducted by Erasmus MC, University Medical Center Rotterdam in close collaboration with the School of Law and Faculty of Social Sciences of the Erasmus University Rotterdam, the Municipal Health Service Rotterdam area, Rotterdam, the Rotterdam Homecare Foundation, Rotterdam and the Stichting Trombosedienst & Artsenlaboratorium Rijnmond (STAR-MDC), Rotterdam. We gratefully acknowledge the contribution of children and parents, general practitioners, hospitals, midwives and pharmacies in Rotterdam. The study protocol was approved by the Medical Ethical Committee of Erasmus MC, Rotterdam. Written informed consent was obtained for all participants. The generation and management of the Illumina 450K methylation array data (EWAS data) for the Generation R Study was executed by the Human Genotyping Facility of the Genetic Laboratory of the Department of Internal Medicine, Erasmus MC, the Netherlands.

The general design of The Generation R Study is made possible by financial support from Erasmus MC, Rotterdam, Erasmus University Rotterdam, the Netherlands Organization for Health Research and Development (ZonMW), and the Ministry of Health, Welfare and Sport. The EWAS data were funded by a grant from the Netherlands Genomics Initiative (NGI)/Netherlands Organisation for Scientific Research (NWO) Netherlands Consortium for Healthy Aging (NCHA; project nr. 050-060-810), by funds from the Genetic Laboratory of the Department of Internal Medicine, Erasmus MC, and by a grant from the National Institute of Child and Human Development (R01HD068437).

We are extremely grateful to all the families who took part in this study, the midwives for their help in recruiting them, and the whole ALSPAC team, which includes data collection staff, data and administrations staff, technical managers and the technical staff with the Bristol Bioresource Laboratory, based within the University of Bristol.

The UK Medical Research Council and Wellcome (Grant ref: MR/Z505924/1) and the University of Bristol provide core support for ALSPAC. Genomewide genotyping data was generated by Sample Logistics and Genotyping Facilities at Wellcome Sanger Institute and LabCorp (Laboratory Corporation of America) using support from 23andMe. A comprehensive list of grants funding is available on the ALSPAC website (http://www.bristol.ac.uk/alspac/external/documents/grant-acknowledgements.pdf). This publication is the work of the authors and C.A.M.C. will serve as a guarantor for the ALSPAC-related contents of this paper.

Analysis of the ALSPAC data was funded by the UK Economic and Social Research Council grant (grant number ES/N000498/1). ARIES was funded by the BBSRC (BBI025751/1 and BB/I025263/1). Supplementary funding to generate DNA methylation data which are (or will be) included in ARIES has been obtained from the MRC, ESRC, NIH and other sources. ARIES is maintained under the auspices of the MRC Integrative Epidemiology Unit at the University of Bristol (grant numbers MC_UU_00011/4 and MC_UU_00011/5).

## Data Sharing Statement

Scripts are available at https://github.com/rosamulder/PrenatalExposome_Epigenome/. Data from the Generation R Study are available upon reasonable request to the director of the Generation R Study, subject to local, national and European rules and regulations. ALSPAC data access is through a system of managed open access. The ALSPAC access policy (http://www.bristol.ac.uk/media-library/sites/alspac/documents/researchers/data-access/ALSPAC_Access_Policy.pdf) describes the process of accessing the data and samples in detail, and outlines the costs associated with doing so.

